# Host protease activity on bacterial pathogens promotes complement- and antibiotic-directed killing

**DOI:** 10.1101/2021.03.05.434144

**Authors:** Shaorong Chen, Alexandria-Jade Roberts, Felix Lu, Carolyn Cannon, Qing-Ming Qin, Paul de Figueiredo

## Abstract

Our understanding of how the host immune system thwarts bacterial evasive mechanisms remains incomplete. Here, we show that host protease neutrophil elastase acts on *Acinetobacter baumannii* and *Pseudomonas aeruginosa* to destroy factors that prevent serum-associated, complement-directed killing. The protease activity also enhances bacterial susceptibility to antibiotics in sera. These findings implicate a new paradigm where host protease activity on bacteria acts synergistically with the host complement system and antibiotics to defeat bacterial pathogens.

---

The human immune system deploys distinct innate immune mechanisms to thwart bacterial pathogens. These mechanisms include the host complement system and host proteases that are present at sites of bacterial infection. The complement system, a network of proteins in sera that are activated by microbial patterns, provides a first line of immune defense (1). Activation results in the deposition of complement proteins on bacterial surfaces, thereby labeling bacteria for phagocytic uptake and subsequent killing. In addition, complement deposition on bacteria drives the formation of the membrane attack complex on bacterial surfaces, which kills Gram-negative bacteria via pore formation. Host proteases, released by immune cells, also contribute to bacterial killing by compromising the integrity of bacterial cell walls (2). In addition, these proteases can destroy virulence factors and thereby thwart bacterial pathogenesis (3). To survive within the human host, bacteria have evolved systems to circumvent, subvert or evade these innate immune defense mechanisms (4). However, the ways in which the host immune system can overcome these immune- evasive bacterial factors constitute a gap in our understanding. Here, we demonstrate that host protease activity on bacterial cell surfaces can destroy bacterial-associated complement inhibitory activities, thereby rendering resistant bacteria susceptible to complement-directed killing and sensitive to frontline antibiotics. Our studies featured the use of three clinically or agriculturally significant bacterial species: *Pseudomonas aeruginosa* (Pa), *Acinetobacter baumannii* (Ab) and *Brucella melitensis* (Bm). Pa and Ab are opportunistic bacterial pathogens that constitute significant threats to civilian and warfighter personnel, as well as patients with underlying disease, including cystic fibrosis (5, 6). Importantly, a significant proportion of clinical isolates of these pathogens display resistance to killing by normal human serum (HS) (7). Moreover, we analyzed a vaccine strain of Bm, the world’s most prevalent bacterial zoonotic agent that displays complement resistance (8).

To test the hypothesis that protease activity on bacteria confers enhanced sensitivity to complement killing in HS, we used a checkerboard strategy (9) to assess synergistic interactions. Briefly, we coincubated the above-mentioned bacterial strains at 1×10^7^ CFU/mL in the presence of various concentrations (0 ~ 20%) of pooled human complement serum (HS, Innovative Research, ICSER100) and neutrophil elastase (NE, EMD Millipore, 324681) (0 ~ 0.3 U/mL) at 37°C. We then determined bacterial growth by measuring the OD_600_ of the culture using a plate reader (BioTek, Inc., VT, USA) at 16 (Pa and Ab) or 72 (Bm) hr post-inoculation (h.p.i.) to assess the inhibitory and/or synergistic activity of HS and NE on the survival or growth of the tested bacteria. Pa strain PAO1 displayed reduced turbidity in 7.5% HS and did not grow when treated with 0.05 U/mL of NE (**Fig. S1A, B**). In 5% HS and 0.1 U/mL of NE, the strain displayed poor growth; however, the turbidity of the cultures was significantly reduced by 0.3 U/mL of NE (**Fig. 1A; Fig. S1A, B**). Pa strain PA14 was weakly resistant to 2.5% HS and growth of this strain was strongly inhibited when treated with greater than 0.1 U/mL NE at this concentration of HS (**Fig. S1A, C**). To explore the hypothesis that NE targeting of protease-labile components on bacterial cell surfaces promoted complement directed killing, we performed similar experiments using PAO1 strains that harbored mutations in the *Ecotin, Wzz* or *AprI* genes. *Ecotin* and *Wzz* contribute to complement resistance (10, 11). AprI is an inhibitor of AprA that protects PAO1 from complement killing. We found that the strains harboring deletions in *Ecotin* and *Wzz* were sensitive to 5% HS (**Fig. 1B-C**), indicating that both *Ecotin* and *Wzz* genes were required for Pa resistance to complement-directed killing. In addition, the PAO1Δ*AprI* mutant displayed enhanced resistance to complement killing and grew well in 5% HS, but was weakly inhibited in 10% HS (**Fig. 1D**). Ab strain Ab5075 displayed resistance to 5% HS but increased sensitivity when treated with NE from 0.1 to 0.3 U/mL (**Fig. 1E**). When treated with 0.3 U/mL NE, the strain displayed reduced growth in 1.25% HS, strong growth inhibition in 2.5% HS, and no growth in 5% HS (**Fig. 1E**). The Bm vaccine strain Bm16MΔ*yjbR* displayed more resistance to complement killing than Ab5075 and PAO1. Bm16MΔ*yjbR* displayed growth in 15% HS in the presence of NE concentrations of ≤ 0.05 U/mL; however, NE treatment increased bacterial sensitivity to HS. When treated with more than 0.1 U/mL of NE, the growth of the Bm strain was inhibited (**Fig. 1F**). Interestingly, Bm16MΔ*yjbR* grew better in HS (< 10%) than non-HS containing medium (**Fig. 1F**). To verify that a heat-labile proteinaceous component of HS was mediating the observed killing activity, we measured bacterial survival in reaction mixtures that contained heat-treated HS. Briefly, HS was heat- inactivated at 55°C for 0.5 h or 65°C for 1 h and then incubated with PAO1 or Ab5075. Under these conditions, no inhibition of bacterial growth was observed, and the synergistic effect observed with non-heat-killed HS was eliminated (**Fig. 1G, H; Fig. S1E**). Interestingly, NE simultaneous coincubation with HS results in better bacterial growth inhibition than subsequent addition of HS (**Fig. 1I, J**), suggesting that a longer period of NE and HS coincubation yields a better synergistic effect in HS (2.5~5%).

**Figure 1.**
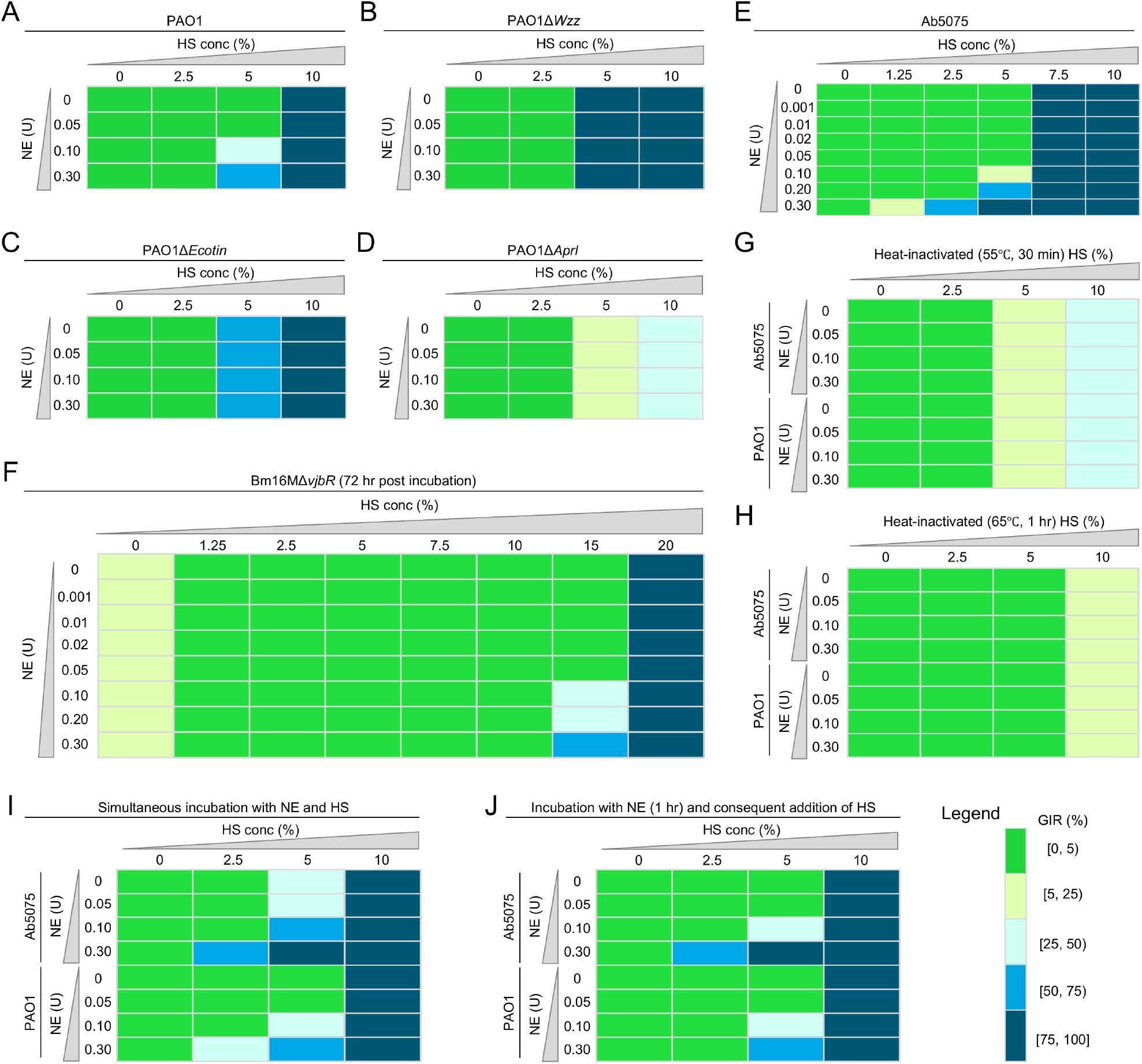
Synergistic effect of neutrophil elastase (NE) and human serum (HS) on bacterial growth. (A-D) Synergy of protease and serum on bacterial pathogens *P. aeruginosa* PAO1 (A) and its derived mutants *PAO1ΔEcotin* (B), PAO1Δ*Wzz* (C), PAO1Δ*AprI* (D). (E, F) NE promotes *A. baumannii* Ab5075 (E) or *B. melitensis* Bm16MΔ*vjbR* (F) killing in the presence of HS. (G, H) Bacterial killing effect of NE decreases in the presence of heat-treated serum at 55°C, 30 min (G) or at 65°C for 1 h (H). (I, J) Addition of NE in HS at the same time (I) or at 1 h post incubation of HS and bacteria (J) varies the synergistic effect. Growth inhibition rate (GIR, %) = [(Contrl OD_600_ – treatment OD_600_)/Contrl OD_600_]×100%. “[” or “]” and “(“ or “)” indicate inclusion and exclusion, respectively.

Taken together, the data support the hypothesis that synergistic interactions between protease activities and complement-directed killing promote the destruction of Gram negative bacterial pathogens. We were intrigued whether these findings could be extended by determining whether protease activity on multi-drug resistant bacteria could enhance sensitivity to front-line antibiotics. Toward this end, we measured the growth of bacteria in the presence of tobramycin (TCI American, Portland, OR, USA). Ab5075 was weakly resistant to 25 μg/mL tobramycin; however, both PAO1 and PA14 displayed sensitivity to tobramycin at this concentration (**Fig. 2A-C**). NE (0.1 to 0.3 U/mL) treated Ab5075 was susceptible to 25 μg/mL tobramycin in 2.5% HS and this treatment displayed a synergistic effect on bacterial killing; NE-treated PAO1 and PA14 were susceptible to 12.5 μg/mL tobramycin (**Fig. 2D-F**). Collectively, these data demonstrated a synergistic interaction between protease activity on bacteria and antibiotic treatment in driving the killing of bacterial pathogens; and providing a new avenue for bacterial disease management.

**Figure 2.**
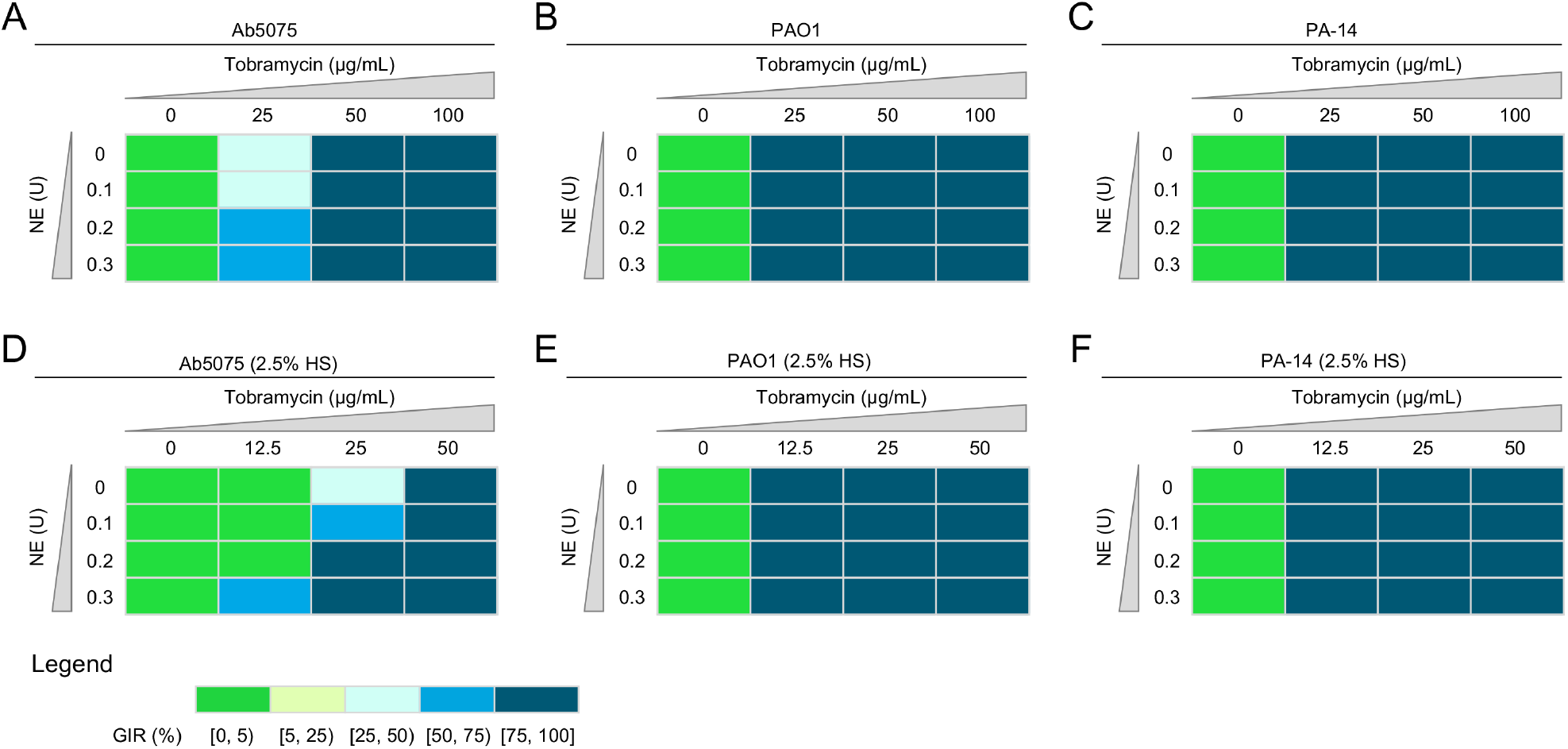
Synergistic effect of NE and tobramycin and/or HS on bacterial killing. (A-C) NE promotes *A. baumannii* Ab5075 (A), *P. aeruginosa* PAO1 (B), or PA-14 (C) killing in the presence of tobramycin. (C-D) Synergy of NE and tobramycin on *A. baumannii* Ab5075 (D), PAO1 (E), or PA-14 (F) killing in the presence of 2.5% HS.

## Data availability

This study did not generate/analyze datasets or code.

## Acknowledgments

This work was supported by the Defense Advanced Research Projects Agency (DARPA) (HR001118A0025-FoF-FP-006), NIH (R21AI139738-01A1, 1 R01AI141607-01A1, 1R21GM132705- 01), the National Science Foundation (DBI 1532188, NSF0854684), the Bill Melinda Gates Foundation and the Texas A&M Clinical Science Translational Research Institute Pilot Grant CSTR2016-1 to PdF. Any opinions, findings, and conclusions or recommendations expressed in this material are those of the author(s) and do not necessarily reflect the views of the funding agencies.

## Supplemental Information

**Figure S1.**
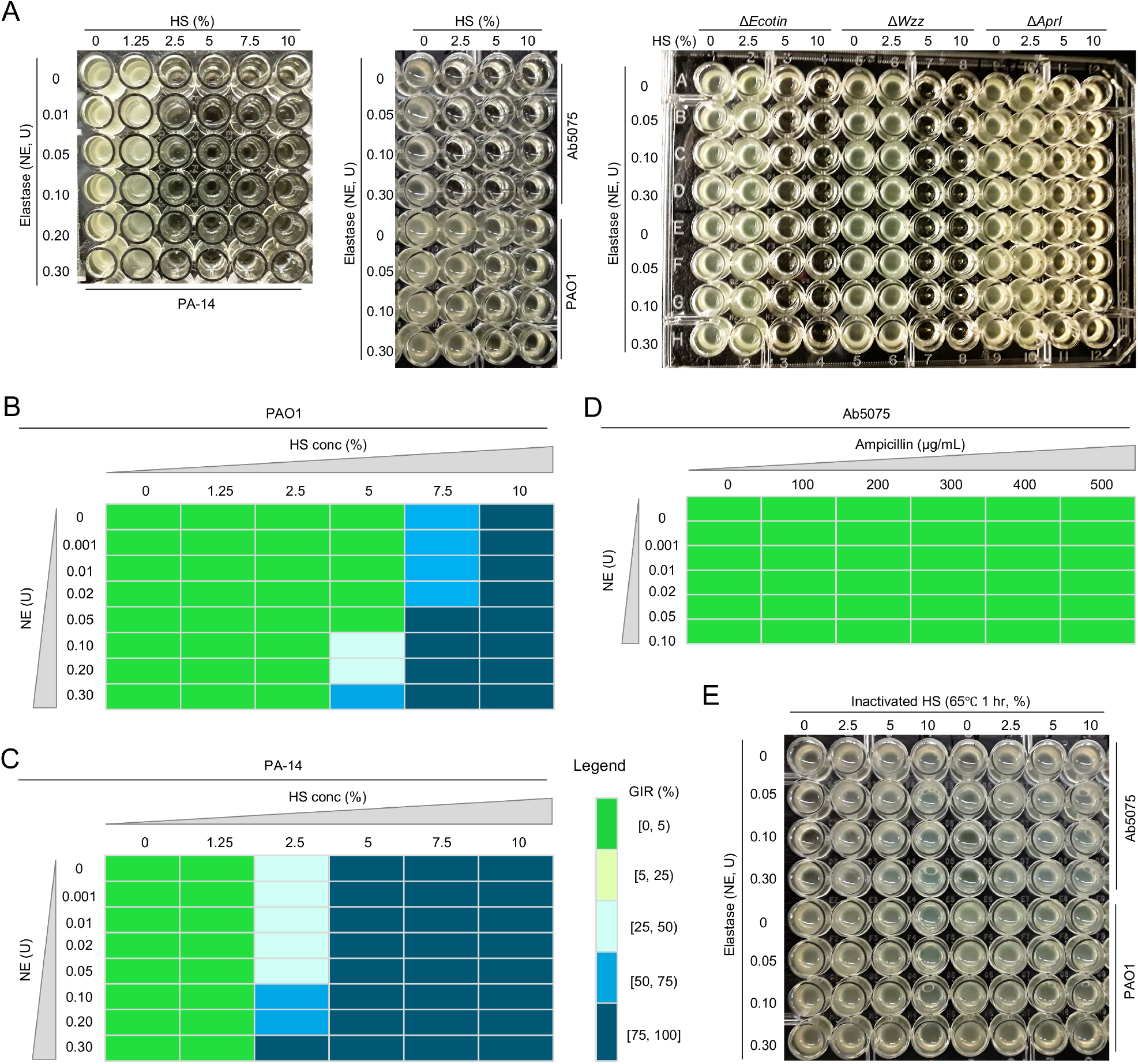
Effect of NE and HS or Ampicillin on bacterial killing. (A) Growth inhibition assays of the bacterial pathogens *P. aeruginosa* PA-14, PAO1, and *A. baumannii* Ab5075 in the indicated concentrations of neutrophil elastase (NE) and normal human serum (HS). (B-C) Synergy of NE and HS on the tested bacterial strains PAO1 (B) and PA-14 (C). (D) NE fails to promote Ab5075 killing in the presence of Ampicillin at the indicated concentrations. (E) Heat-inactived HS fails to promote NE bacterial killing. Growth inhibition rate (GIR, %) = [(Contrl OD_600_ – treatment OD_600_)/Contrl OD_600_]×100%. “[” or “]” and “(“ or “)” indicate inclusion and exclusion, respectively. Pictures froms a representative experiments of at least three independent experiments.

## Notes

### Competing Interest Statement

The authors have declared no competing interest.

### Summary of Updates

change author list

## References

1. Heesterbeek DAC, Angelier ML, Harrison RA, Rooijakkers SHM. Complement and Bacterial Infections: From Molecular Mechanisms to Therapeutic Applications. J Innate Immun. 2018;10(5-6):455–64.

2. Tarhini M FH, Bentaher A, Greige-Gerges H, Elaissari A. A potential new strategy for using elastase and its inhibitor as therapeutic agents. Journal of Translational Science. 2018.

3. Kobayashi SD, Malachowa N, DeLeo FR. Neutrophils and Bacterial Immune Evasion. J Innate Immun. 2018;10(5-6):432–41.

4. Janeway Jr CA, Travers P, Walport M, Shlomchik MJ. Pathogens have evolved various means of evading or subverting normal host defenses. Immunobiology: The Immune System in Health and Disease 5th edition: Garland Science; 2001.

5. Diggle SP, Whiteley M. Microbe Profile: Pseudomonas aeruginosa: opportunistic pathogen and lab rat. Microbiology (Reading). 2020;166(1):30–3.

6. Howard A, O’Donoghue M, Feeney A, Sleator RD. Acinetobacter baumannii: an emerging opportunistic pathogen. Virulence. 2012;3(3):243–50.

7. Sanchez-Larrayoz AF, Elhosseiny NM, Chevrette MG, Fu Y, Giunta P, Spallanzani RG, et al. Complexity of Complement Resistance Factors Expressed by Acinetobacter baumannii Needed for Survival in Human Serum. J Immunol. 2017;199(8):2803–14.

8. Hensel ME, Garcia-Gonzalez DG, Chaki SP, Hartwig A, Gordy PW, Bowen R, et al. Vaccine Candidate Brucella melitensis 16MΔvjbR Is Safe in a Pregnant Sheep Model and Confers Protection. Msphere. 2020;5(3).

9. Martinez-Irujo JJ, Villahermosa ML, Alberdi E, Santiago E. A checkerboard method to evaluate interactions between drugs. Biochemical pharmacology. 1996;51(5):635–44.

10. Kintz E, Scarff JM, DiGiandomenico A, Goldberg JB. Lipopolysaccharide O-antigen chain length regulation in Pseudomonas aeruginosa serogroup O11 strain PA103. J Bacteriol. 2008;190(8):2709–16.

11. Tseng BS, Reichhardt C, Merrihew GE, Araujo-Hernandez SA, Harrison JJ, MacCoss MJ, et al. A Biofilm Matrix-Associated Protease Inhibitor Protects Pseudomonas aeruginosa from Proteolytic Attack. mBio. 2018;9(2).

